# pNeRF: Parallelized Conversion from Internal to Cartesian Coordinates

**DOI:** 10.1101/385450

**Authors:** Mohammed AlQuraishi

## Abstract

The conversion of polymer parameterization from internal coordinates (bond lengths, angles, and torsions) to Cartesian coordinates is a fundamental task in molecular modeling, often performed using the Natural Extension Reference Frame (NeRF) algorithm. NeRF can be parallelized to process multiple polymers simultaneously, but is not parallelizable along the length of a single polymer. A mathematically equivalent algorithm, pNeRF, has been derived that is parallelizable along a polymer’s length. Empirical analysis demonstrates an order-of-magnitude speed up using modern GPUs and CPUs. In machine learning-based workflows, in which partial derivatives are backpropagated through NeRF equations and neural network primitives, switching to pNeRF can reduce the fractional computational cost of coordinate conversion from over two-thirds to around 10%. An optimized TensorFlow-based implementation of pNeRF is available on GitHub.

## Introduction

Polymers can be mathematically represented by the Cartesian coordinates of their atoms, or by a sequence of bond lengths, angles, and torsions of adjacent bonded atoms (internal coordinates).^[1]^ Each parameterization has its own advantages and disadvantages. In Cartesian space, spatially proximal atoms that are distant along the polymer chain can be readily detected, facilitating distance-based computations involving e.g. electrostatics. When sampling changes in polymer conformations however, the Cartesian parameterization can be brittle, leading to non-physical clashes which the internal coordinates parameterization avoids by explicitly modeling bonded interactions.^[2]^ Consequently, rapid interchanging between the two parameterizations is critical for many established molecular modeling applications, including molecular dynamics and Monte Carlo-based sampling.^[3]^ Certain force fields, such as the Rosetta^[4]^ energy function for biomolecules, explicitly encode Cartesian and internal energy terms and therefore require simultaneous use of both parameterizations. Emerging applications using machine learning-based (ML) molecular modeling, in which force fields^[5], [6]^ or molecules^[7]^ are optimized by backpropagating partial derivatives through the internal and Cartesian coordinates of polymers, further necessitate computing the derivatives of the internal-to-Cartesian transformation equations.

A widely used method for performing this computationally-demanding transformation is the Natural Extension Reference Frame (NeRF) algorithm.^[1]^ When transforming multiple independent chains, NeRF is embarrassingly parallelizable, linearly scaling in parallelization capacity with the number of polymers. However for a single polymer, NeRF runs sequentially. While not a bottleneck for CPUs with limited core counts, modern CPUs and GPUs provide massive parallelism that is seldom saturated by NeRF in conventional molecular modeling pipelines. Additionally for ML-based workflows, there are limits on the number of polymers that can be processed simultaneously, as the generalization quality of learned models is often inversely related to the batch size (number of data points used to estimate the gradient) used in training them.^[8]^ Combined with the fact that ML workflows perform a large number of evaluations during model training, the computational cost of NeRF evaluations can be substantial.

We derive a new algorithm, pNeRF, that is mathematically equivalent to NeRF but is parallelizable even for a single polymer chain, with a total computational cost equal to NeRF plus *M* additional affine transformations, where *M* is the number of parallel threads used. We empirically show that on both modern CPUs and GPUs, pNeRF can be over an order-of-magnitude faster than NeRF. We further demonstrate that on realistic ML-based workflows, use of pNeRF reduces the fractional cost of internal-to-Cartesian coordinate conversion from 67% to 13%. Finally we provide an empirical analysis of optimal usage criteria based on polymer lengths, number of independent polymers processed, and CPU vs. GPU parallelism.

## Methods

### NeRF

We begin with a summary of the standard NeRF algorithm. Given a sequence of bond lengths, angles, and torsions of adjacent bonded atoms, NeRF sequentially constructs a linear polymer from one end of the molecule to the other (extensions for branched polymers are straightforward.) First the coordinates of each atom, encoded by a triplet of length, angle, and torsion, are computed in a special reference frame (SRF), possibly in parallel. The algorithm then sequentially moves each atom from the SRF to its actual position using an affine transformation derived from the coordinates of the three previously computed atoms. Formally, let *r, θ, φ* be the bond length, angle, and torsion of an atom with respect to its preceding neighbors, then

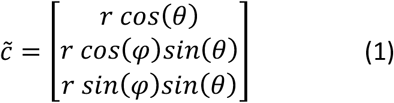

is its SRF coordinates. Given a previously computed sequence of coordinates *c*_2_, … , *c*_*k*−1_, the next set of coordinates is 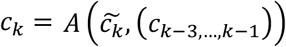 where 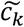 is the SRF set of coordinates and 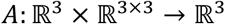 is a function mapping the SRF coordinates to the actual position using a rigid transformation determined by the last three coordinates. Specifically, letting *m_k_* = *c*_*k*−1_ – *c*_*k*−2_ and 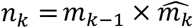 where 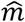 is the unit-normalized version of *m* and × is the cross product, then

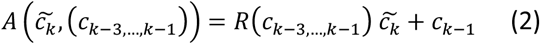

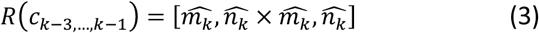

where 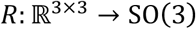 is a function mapping the previous three coordinates to a rotation matrix.^[1]^ By sequentially applying *A* to the sequence of SRF coordinates 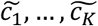 of a length *K* polymer, NeRF converts internal coordinates into Cartesian coordinates. The choice of the initial three coordinates used to transform 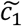 is arbitrary, which we term the initialization coordinates.

### pNeRF

Because NeRF requires the coordinates of the last three atoms to position the next atom, it does not permit parallelization along the polymer length. The basic idea behind pNeRF is to fragment the polymer into *M* equal-sized fragments, independently convert each into Cartesian space, and then reassemble the fragments into the final polymer (Figure 1). A naïve implementation of this idea would involve four affine transformations and a matrix inversion for each of the *M* fragments. We derive a formulation that adds only one affine transformation per fragment.

**Figure 1:**
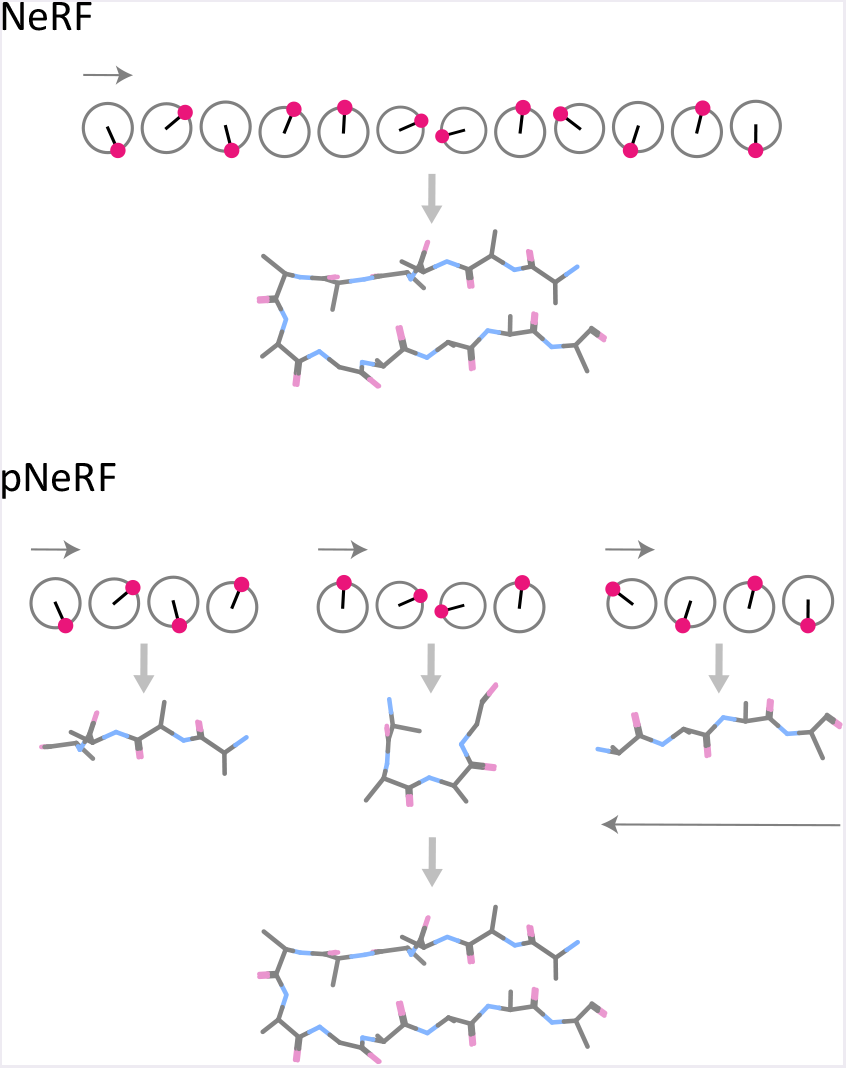
In the standard NeRF algorithm, internal coordinates (angles and bond lengths, shown as dots on a circle) are converted to Cartesian coordinates (shown as sticks) sequentially, starting from one end of the polymer and finishing at the opposite end. In pNeRF, multiple fragments are reconstructed independently and in parallel, and then the final coordinates are obtained by reorienting entire fragments, sequentially, in the opposite direction.

### Algorithm

Let 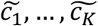 be the SRF coordinates of a polymer of length *K*, and without loss of generality assume that *M* divides *K*. Partition the coordinates into *M* subsets 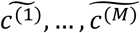 such that 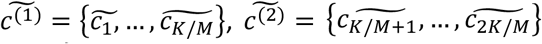, … Using these initialization coordinates (columns are coordinates):

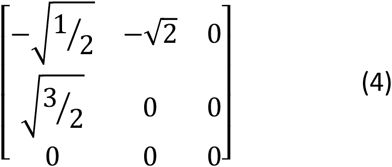

apply NeRF independently to each subset. Once complete, apply 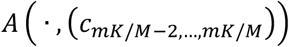 to every entry in 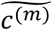 for *m* = 1, …, *M*, then concatenate the subsets. The resulting sequence is equivalent (up to a rigid transformation) to a sequential coordinate conversion using NeRF.

### Proof of correctness

#### Proposition 1

Given initialization coordinates ***x*** = (*x*_1_, *x*_2_, *x*_3_), applied sequentially to a sequence of SRF coordinates 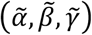 (e.g. the beginning of a new fragment), i.e.

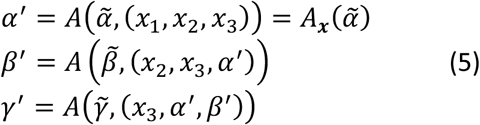

and a previously computed sequence of coordinates ***c*** = {…, *c*_*k*−2_, *c*_*k*−1_, *c_k_*] used to transform 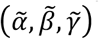 to their final location, i.e.

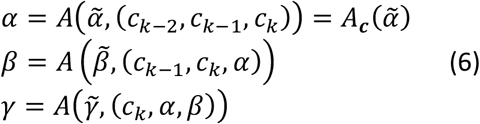

the following relationships hold true:

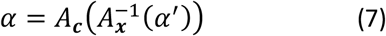

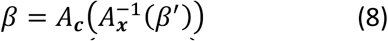

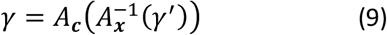

where *A*^−1^(*δ*, (*α, β, γ*)) = *R*^−1^(*α, β, γ*) (*δ − γ*) and *R*^−1^ is always defined as *R* is a rotation matrix. We abbreviate *A*(*η*, (*x*_1_,*x*_2_,*x*_3_)) and *A*(*η*, (*c*_*k*−2_,*c*_*k*−1_,*c_k_*)) using *A_x_* and *A_c_*, respectively, and similarly for *R_x_* and *R_c_*.

Note that 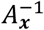 is fixed and independent of the coordinates, and hence can be pre-computed. Note also that since 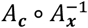 is a rigid affine transformation that brings (*α′, β′, γ′*) into alignment with (*α,β,γ*), then for an arbitrary new coordinate 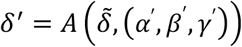, the following must hold true:

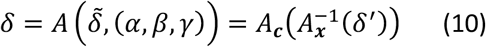

and by induction, all subsequent coordinates must be similarly transformed. Thus if proposition 1 is true, we can independently compute *M* fragments *c*^(1)^, … , *c*^(*M*)^ and sequentially transform them using 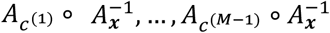 to their final correct positions. We can further simplify the procedure by choosing (*x*_1_, *x*_2_, *x*_3_) such that 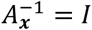 (this computation is provided after the proof).

Before we proceed with the proof, we first introduce a lemma and some corollaries.

#### Lemma 1

Let *R* be a function as defined in equation 3 and *R′* be any rotation matrix, then

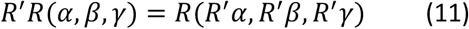

To prove this we will consider each column vector of *R*(*α, β, γ*) separately, starting with the first column:

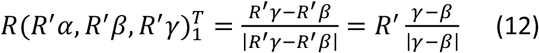

where we used the fact that rotations do not alter vector magnitude. For the third column:

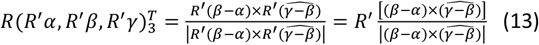

where we used the fact that rotations are distributive over cross products. The same arguments apply for the second column, and thus we obtain the lemma.

#### Corollary 1

Let *A* be a function as defined in equation 2, then

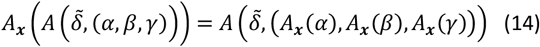

To prove this corollary we work from the right-hand side:

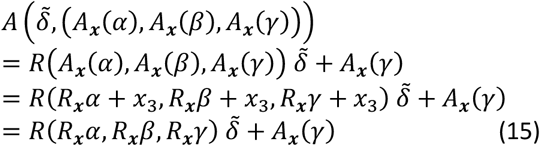

where the last step used the fact that by construction, *R* is invariant to a translation of its arguments. Applying lemma 1, we obtain:

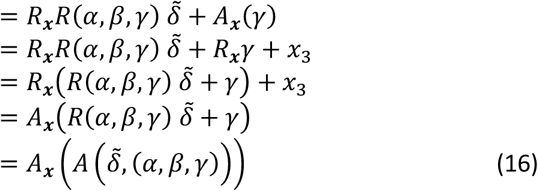

Note that (14) holds for inverses as well, i.e.

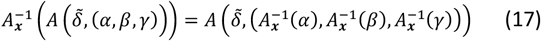

since 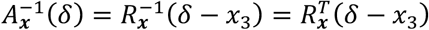 as 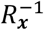 is a rotation matrix and thus lemma 1 is applicable to 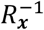.

#### Corollary 2

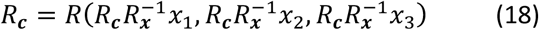

For any 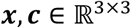. The result follows from applying lemma 1 to 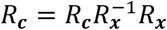.

We are now ready to prove proposition 1.

#### Proof for *α* (eq. 7)

Trivially follows from definitions (eqs. 5 and 6)

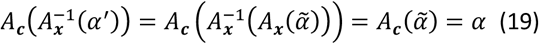

#### Proof for *β* (eq. 8)

By definition (eq. 5):

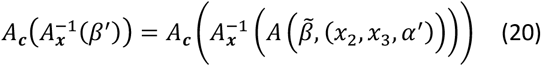

Applying corollary 1 to innermost *A* in rhs we get:

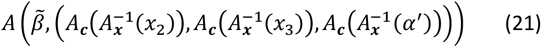

From before 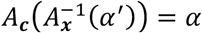. We will work out the other arguments to *A*:

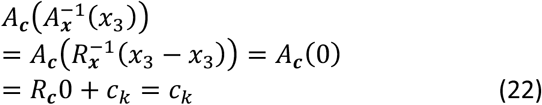

Similarly we have (applying corollary 2 in the last step since eq. 23 is an argument to *A*):

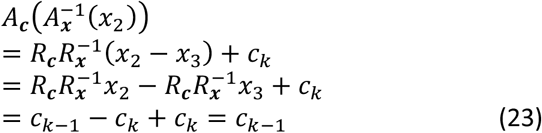

This implies that eq. 21 is equal to 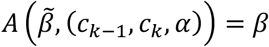 which proves eq. 8.

#### Proof for *γ* (eq. 9)

Starting with the definitions and applying corollary 1 as we did for eq. 8 we obtain

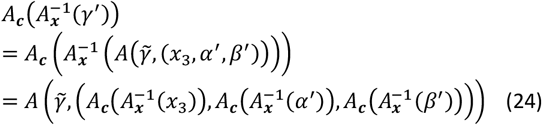

Applying eqs. 22, 19, and 20 to the first, second, and third arguments of eq. 24, respectively, we get 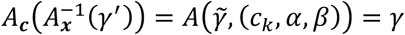. This proves proposition 1.

### Initialization coordinates

In eqs. 7-9 the initialization coordinates ***x*** = (*x*_1_, *x*_2_, *x*_3_) can be arbitrarily chosen. A judicious choice of ***x*** can yield 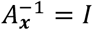, eliminating one extraneous affine transformation per fragment. We derive one such set of coordinates next.

First note that 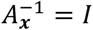 if and only if *α* = *A_**c**_*(*α*′). We start with eq. 6 and apply the above condition followed by corollary 1:

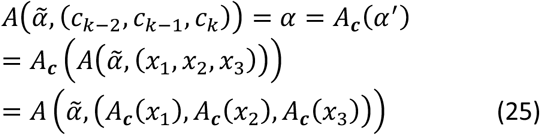

This yields the following set of equations:

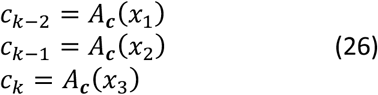

which we solve to obtain the desired ***x***. For *x*_3_ we obtain a unique solution:

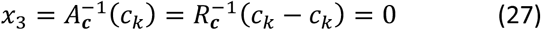

consistent with the fact that for an affine transformation to be the identity its translation component must be 0.

For *x*_2_, we left-multiply by an arbitrary *R* of our choosing, then apply corollary 2 to obtain:

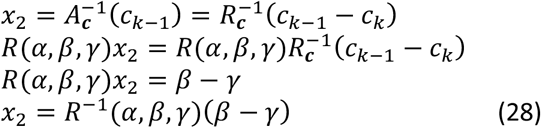

The above provides an explicit solution for *x*_2_, and similarly for *x*_1_:

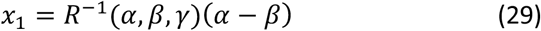

Since the choice of *α, β, γ* is arbitrary, we choose the standard basis, and obtain eq. 4 as the solution.

## Results and Discussion

### pNeRF is up to 13x faster than NeRF

We implemented pNeRF using the TensorFlow^[9]^ automatic differentiation^[10]^ framework. This enables its use in both conventional applications in which internal coordinates are simply converted to Cartesian coordinates (“forward pass”), and machine learning-based applications in which the derivatives of such conversions are backpropagated from a loss function to the parameters of a learned model (“backward pass”).^[11]^ Using the TensorFlow implementation, we assessed pNeRF’s performance on realistic settings—in terms of sequence lengths and batch sizes (number of simultaneous conversions)—using modern CPUs (Xeon E5-2643 v4) and GPUs (Titan Xp). We considered batch sizes ranging in size from 1 to 512 in doubling increments, and sequence lengths ranging from 100 to 1,000 in increments of 100. For each combination of batch size and sequence length, we carried out the forward and backward passes of pNeRF 110 times, and averaged the timings of the last 100 passes (the first 10 are used to burn-in the process and minimize variability.) Experiments were done using 1, 5, 15, and 25 fragments (M), and the fastest option was chosen for each batch size / sequence length combination.

Figure 2 shows the timings, where intensities correspond to fold speed up resulting from using pNeRF over NeRF. In general the same trends can be observed for CPUs and GPUs, and the forward and backward passes, with longer sequences and smaller batch sizes gaining more from pNeRF than shorter sequences and larger batch sizes. This is expected as longer sequences enable greater parallelism, while larger batch sizes saturate the computational throughput of CPUs and GPUs. We observe speed ups of up to 13x in the configurations we considered, although in principle the speed up is not bounded, and future processors with greater capacity for parallelism will yield even larger benefits. We never observed slowdowns due to excessive parallelization by pNeRF.

**Figure 2.**
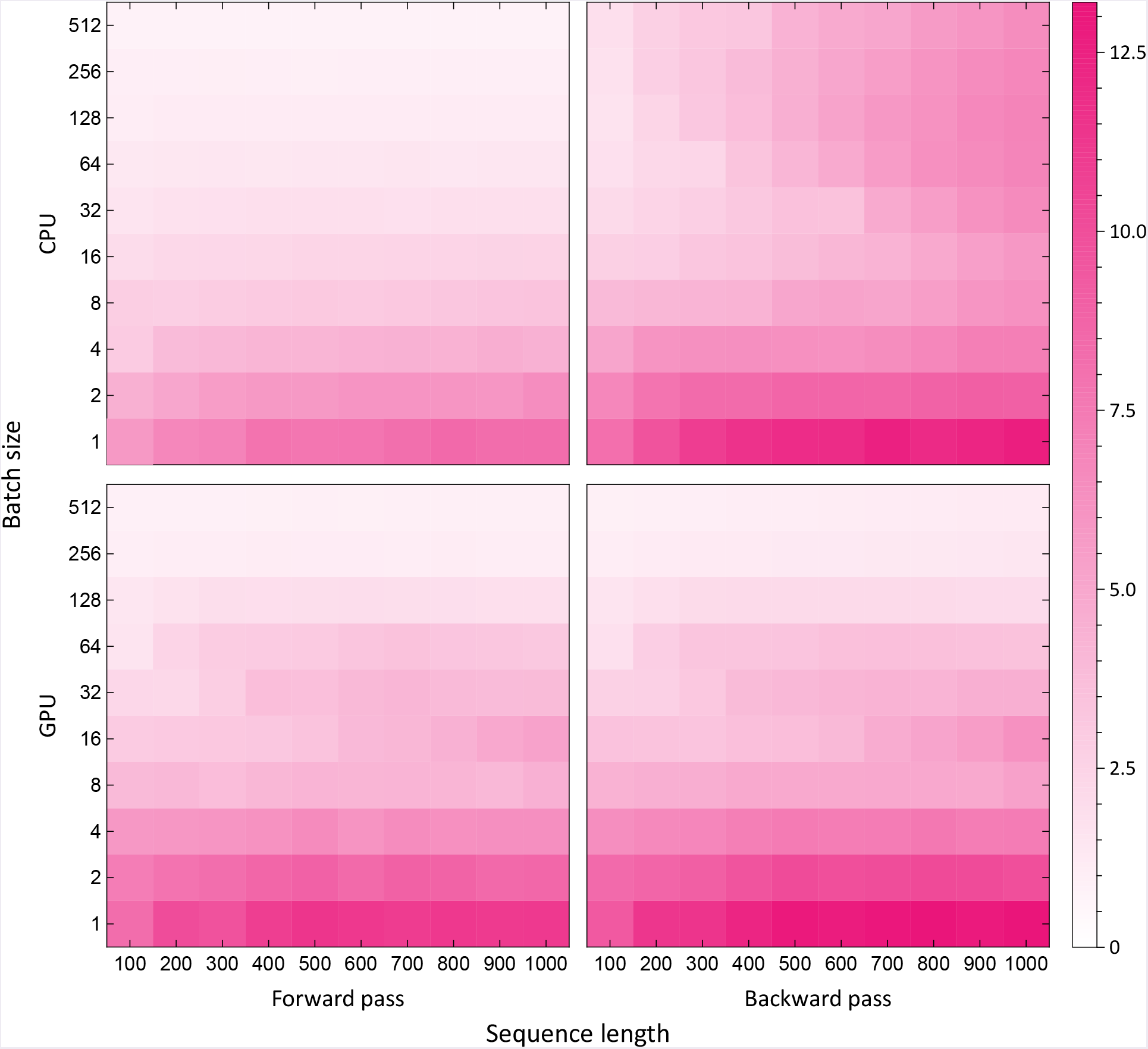
Fold speed up in computation time (pink intensity) when using pNeRF instead of NeRF for different combinations of batch sizes and sequence lengths. Computations were carried out on CPUs (Xeon E5-2643 v4) and GPUs (Titan Xp) and represent the averages of 100 independent runs, preceded by 10 burn-in runs.

### Optimal hardware choice is model-dependent

pNeRF relies heavily on trigonometric operations, which do not necessarily exploit the computing capabilities of GPUs, particularly if the opportunities for parallelism are limited (e.g. short sequences.) This suggests that the choice of optimal hardware may depend on batch size and sequence length. To assess this, we computed the log ratio of processing times on CPUs versus GPUs, shown in Figure 3. Values above 0 correspond to configurations were GPUs are faster, and values below 0 indicate CPUs are faster. In general, we observe that GPUs outperform CPUs for batch sizes of 64 and larger, if the sequences are at least 200 – 300 steps long. For smaller batch sizes, CPUs dominate irrespective of sequence length. We also observe that for very large batch sizes during the forward pass, memory limitations on GPUs can result in poor performance relative to CPUs.

**Figure 3:**
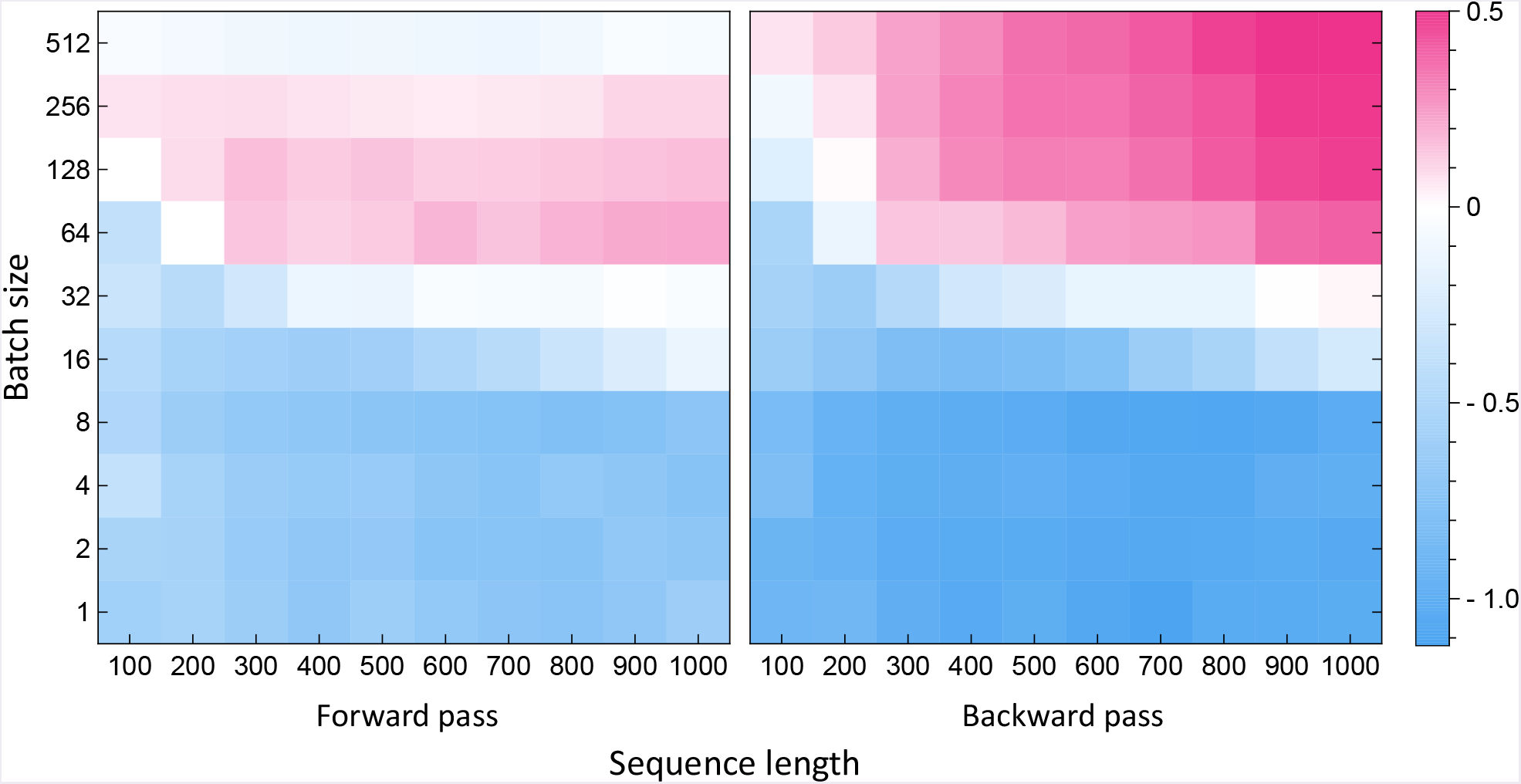
Log ratio of pNeRF CPU over GPU compute time (>0 indicate GPUs are faster) for different combinations of batch sizes (number of simultaneous conversions) and sequence lengths.

### Optimal number of fragments is model- and hardware-dependent

pNeRF introduces a free parameter, *M*, which controls the number of fragments converted in parallel. To obtain maximum throughput, this parameter must be optimized for the given choice of batch size, sequence length, and hardware. Figure 4 illustrates pNeRF’s behavior for varying batch sizes and hardware platforms, assuming a fixed sequence length of 700 steps, as a function of *M*. Arrows indicate the best performing choice of *M* for each configuration. Numbers were computed in the same way as in Fig. Figure 2 and Figure 3. Not surprisingly, smaller batch sizes permit higher numbers of fragments, as the processor is not yet saturated. Furthermore, the choice of optimal processor (and associated *M*) changes depending on the batch size, with CPUs performing best for batches of size 1 and 8 and GPUs for batches of size 64 and 512. In general we see agreement between the forward and backward passes, simplifying implementation.

**Figure 4:**
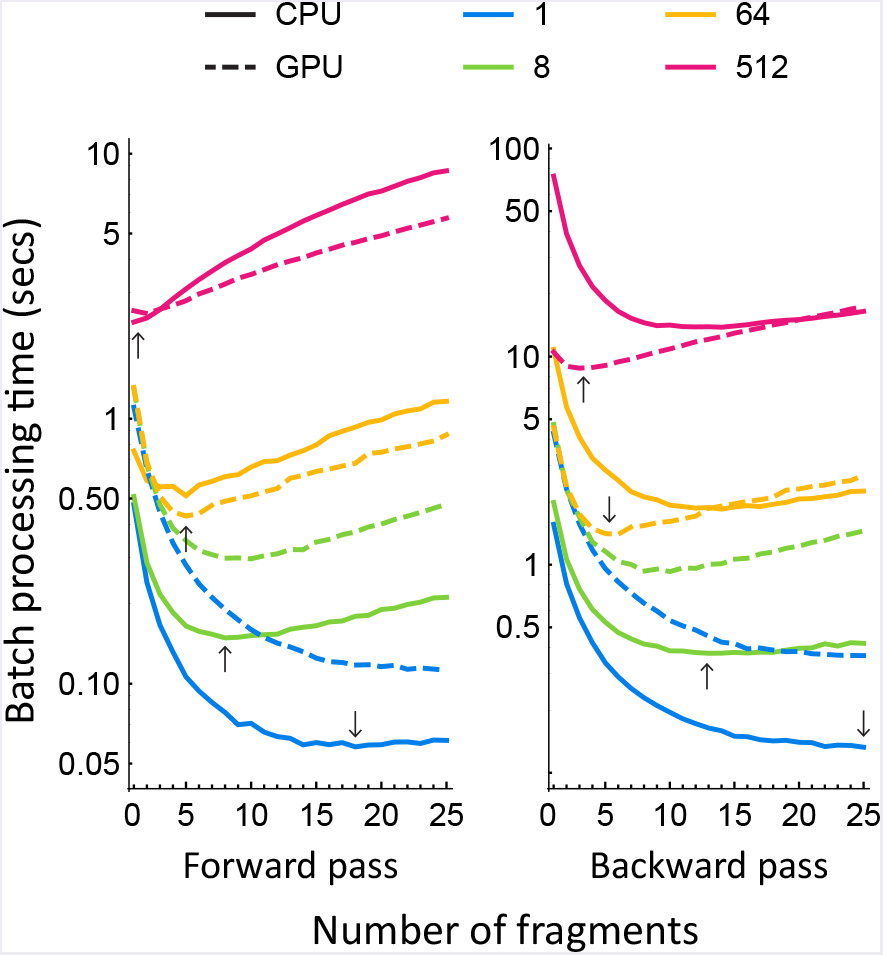
pNeRF processing times for different batch sizes (indicated by numbers in legend) and hardware platforms as a function of the number of fragments (M) used by pNeRF in the forward and backward passes. Arrows indicate the fastest choice of *M* for each configuration. All runs used a sequence length of 700. Timings represent the averages of 100 independent runs, preceded by 10 burn-in runs.

### pNeRF performance in machine learning-based workflows

In practical applications the conversion between internal and Cartesian coordinates is not done in isolation but is instead part of a larger workflow. We sought to assess the impact of switching from NeRF to pNeRF in a real-world machine learning model that utilizes the forward and backward passes of pNeRF computations. Recurrent geometric networks^[12]^ (RGNs), which differentiably learn a mapping from protein sequence to structure, are one such model. They integrate trainable computations, known as Long Short-Term Memory^[13]^ (LSTM), with geometric operations including the conversion from internal to Cartesian coordinates. We assessed the batch processing time for different variants of the RGN architecture, using both NeRF and pNeRF.

Figure 5 shows the results for two choices of architectures (top line on x-axis, denoting number of LSTM layers x layer size) and maximum sequence lengths (bottom line on x-axis.) The LSTM contribution to compute time is shown in blue, while the (p)NeRF contribution is in pink. Left (pink) bars correspond to standard NeRF, and right bars are to pNeRF. All timings shown are for batch sizes of 32, which were comprised of real data from the Protein Data Bank.^[14]^ We observe that while NeRF can account for a major portion of total RGN compute time, around 2/3 for the smaller LSTM architecture, it is reduced to a negligible level (~10%) when using pNeRF. This demonstrates practical utility in an emerging application, and it is likely that future workflows making more extensive use of pNeRF will see greater gains.

**Figure 5:**
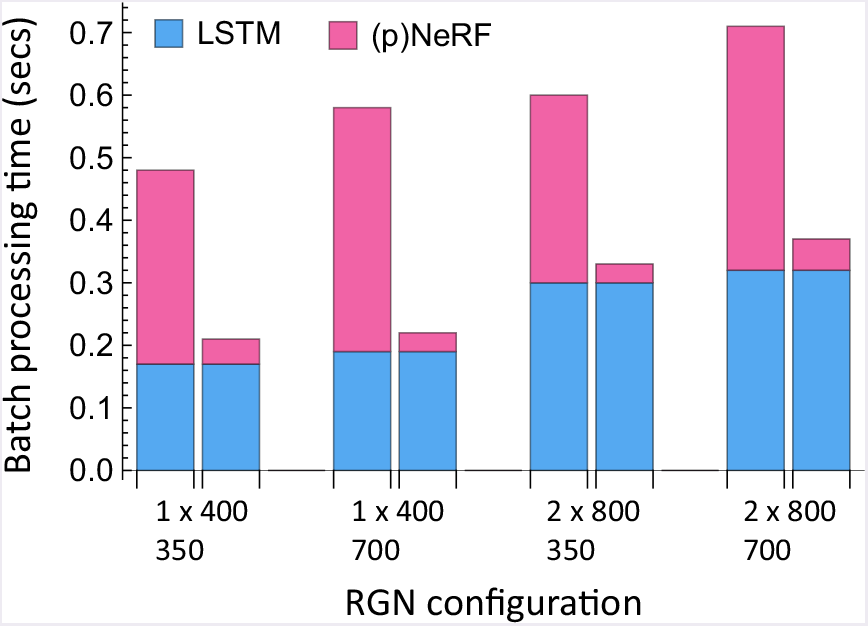
Contribution to RGN batch processing time from LSTM and (p)NeRF components, computed using different choices of architectures and maximum sequence lengths. NeRF contributions are shown in the left bars, and pNeRF contributions are shown in the right bars. The first line of RGN configuration corresponds to number of bidirectional LSTM layers x layer size, while the second line indicates maximum sequence length.

Note that the lack of a major timing difference between maximum sequence lengths is due to the relatively short average sequence length of proteins in the PDB (~300), which limits computational cost as longer sequences are not frequently encountered.

## Conclusions

We derived pNeRF, a mathematically equivalent algorithm to NeRF that enables virtually unbounded parallelism subject to hardware restrictions. We characterized its behavior under different experimental conditions and showed that it can lead to substantial speed gains on real-world applications. While the use of geometric transformations—including internal-to-Cartesian coordinate conversion—in machine learning applications is in its nascent stage, the rapid growth of deep learning models in the molecular sciences will likely lead to increased use of such transformations. Consequently we believe that pNeRF will find broad use in a variety of applications, particularly in polymer science. To facilitate further use and development of pNeRF we have made public a high-performance TensorFlow-based implementation, suitable for machine learning applications, on GitHub at https://github.com/aqlaboratory/pnerf.

## Acknowledgments

We thank Peter Sorger for his mentorship and support, and Uraib Aboudi for her feedback. We gratefully acknowledge the support of NVIDIA Corporation with the donation of the Titan Xp GPUs used for this research. This work was supported by NIGMS Grant P50GM107618.

## References and Notes

1 J. Parsons, J.B. Holmes, J.M. Rojas, J. Tsai, C. E. M. Strauss, J. Comput. Chem., 2005, DOI:10.1002/jcc.20237.

2 N. Vaidehi, A. Jain, J. Phys. Chem. B, 2015, D0I:10.1021/jp509136y.

3 D. Marx, J. Hutter, in Ab initio molecular dynamics: basic theory and advanced methods; Cambridge University Press, Cambridge, 2012.

4 A. Leaver-Fay, M. Tyka, S. M. Lewis, O. F. Lange, J. Thompson, R. Jacak, K. Kaufman, P. D. Renfrew, C.A. Smith, W. Sheffler, I. W. Davis, S. Cooper, A. Treuille, D. J. Mandell, F. Richter, Y.-E. A. Ban, S. J. Fleishman, J. E. Corn, D. E. Kim, S. Lyskov, M. Berrondo, S. Mentzer, Z. Popović, J. J. Havranek, J. Karanicolas, R. Das, J. Meiler, T. Kortemme, J. J. Gray, B. Kuhlman, D. Baker, P. Bradley, Meth. Enzymol., 2011, DOI:10.1016/B978-0-12-381270-4.00019-6.

5 J. S. Smith, O. Isayev, A. E. Roitberg, Chem. Sci., 2017, DOI:10.1039/C6SC05720A.

6 J. M. Jumper, K. F. Freed, T. R. Sosnick, bioRxiv, 2017, DOI:10.1101/169326.

7 M. Ragoza, L. Turner, D. R. Koes, arXiv:1710.07400[cs, q-bio, stat], 2017.

8 D. Masters, C. Luschi, arXiv:1804.07612 [cs, stat], 2018.

9 Martín Abadi, A. Agarwal, P. Barham, E. Brevdo, Z. Chen, C. Citro, G. S. Corrado, A. Davis, J. Dean, M. Devin, S. Ghemawat, I. Goodfellow, A. Harp, G. Irving, M. Isard, Y. Jia, R. Jozefowicz, L. Kaiser, M. Kudlur, J. Levenberg, D. Mané, R. Monga, S. Moore, D. Murray, C. Olah, M. Schuster, J. Shlens, B. Steiner, I. Sutskever, K. Talwar, P. Tucker, V. Vanhoucke, V. Vasudevan, F. Viégas, O. Vinyals, P. Warden, M. Wattenberg, M. Wicke, Y. Yu, X. Zheng, TensorFlow: Large-Scale Machine Learning on Heterogeneous Systems. 2015.

10 A. G. Baydin, B. A. Pearlmutter, A. A. Radul, J. M. Siskind, arXiv:1502.05767[cs], 2015.

11 I. Goodfellow, Y. Bengio, A. Courville, in Deep Learning; The MIT Press, Cambridge, Massachusetts, 2016.

12 M. AlQuraishi, bioRxiv, 2018, DOI:10.1101/265231.

13 S. Hochreiter, J. Schmidhuber, Neural Computation, 1997, DOI:10.1162/neco.1997.9.8.1735.

14 F. C. Bernstein, T. F. Koetzle, G. J. Williams, E. F. Meyer, M. D. Brice, J. R. Rodgers, O. Kennard, T. Shimanouchi, M. Tasumi, J.Mol. Biol., 1977, 112, 535–542.

